# Cue-Dependent Fear Learning Drives Nucleus Accumbens Spine Plasticity

**DOI:** 10.64898/2026.02.25.707962

**Authors:** Desh Deepak Ratna, Cortez Gray, Eugene Lee, Harris Kiaris, Makenna Hamilton, T. Chase Francis

## Abstract

Nucleus accumbens (NAc) dopamine 2 receptor expressing medium spiny neurons (D2-MSNs) are involved in stress and aversive learning, where repeated stress increases excitatory spine density. Whether this plasticity reflects cue-specific learning or generalized stress response remains unknown. Using Pavlovian fear conditioning in Tac1-Cre/Tdtomato mice, we dissociated associative plasticity from the effects of foot shock stress. Acute fear conditioning produced distinct physiological outcomes between stress in the presence or absence of a cue. Conditioning for 7 days consolidated cue learning and increased excitatory transmission frequency via an increase in the total spine density. However, repeated exposure to foot shock did not lead to this synaptic remodeling. Our results suggest that morphological changes supporting synaptic plasticity on NAc D2-MSNs are due to cue-dependent learning, but not foot shock stress alone. We propose that NAc D2-MSNs encode learning and response to threat cues, which may heighten later stress responsivity.

## Introduction

Stress-related brain areas detect and encode environmental cues based on their valence and salience, which helps organisms distinguish between safe and threatening conditions, and is required for survival. Acute stress is adaptive and can mobilize energy reserves for a reactive response and enhance aversive learning to threatening stimuli ^1–3^. However, prolonged stress exposure causes maladaptive alterations in neuronal structure and signaling ^4,5^, resulting in pathological conditions including depression and anxiety ^6,7^. However, the number and duration of stressors required to induce these structural and physiological changes, as well as the intersection between learning and negative emotional outcomes, have not been thoroughly explored.

The Nucleus Accumbens (NAc) has recently been identified as a key locus necessary for the expression of stress and aversive responses ^8,9^. NAc medium spiny projection neurons (MSNs), characterized by their dopamine 1 (D1) or dopamine 2 (D2) receptor expression, play distinct roles in processing aversive stimuli, where D1-MSNs are required for cue-dependent aversive conditioning, and D2-MSNs are necessary for cue-dependent recall ^10–12^. In addition, D2-MSNs have been implicated in learning in other contexts, including negative reinforcement ^13,14^, decision-making ^12,15^, reversal learning ^16,17^, and reward learning ^12,14^. These studies indicate that D2-MSNs play a more subtle role in aversive response and may not directly contribute to negative outcomes to stress.

NAc MSN subtypes display dichotomous excitatory synaptic changes following repeated stress. Interestingly, despite depression-related outcomes being attributed to decreased NAc function ^8,18,19^, repeated stress increases the density of excitatory spines on D2-MSNs and increases the frequency of excitatory input ^8,9,20,21^. Whereas in D1-MSNs, structural changes after repeated stress reduce excitatory transmission and D1-MSN activity bidirectionally alters depression-related outcomes ^5,8,22–25^. Further evidence indicates that activating D2-MSNs during an acute, mild social stressor promotes social avoidance without affecting hedonic outcomes ^8^, and spine density correlates with social avoidance ^21^. Regardless of the animal’s stress-susceptibility state, these observations suggest that D2-MSNs facilitate a learned or generalized avoidance response towards a stressor or aversive stimuli, helping identify harmful environmental cues. This study aims to determine how NAc D2-MSN morphological plasticity confers specific cued features about stressful stimuli.

In this study, we tested the hypothesis that plasticity in NAc D2-MSNs is driven by conditioned aversive cues rather than by repeated non-conditioned foot shock. Pavlovian fear conditioning was used to assess aversive learning and to provide a repeated foot shock stressor. Surprisingly, we found plasticity on NAc D2-MSNs shifted across repeated foot shock from an increase in amplitude to an increase in the frequency of excitatory synaptic input. Our study further provided proof of principle by testing how substance P-dependent signaling, which is necessary for NAc D2-MSN plasticity and cue-dependent learning, blunts this plasticity and learning through an unexpected decrease in release probability. Our data underscores the importance of NAc D2-MSN excitatory plasticity in learning about aversive stimuli and diminishes the role of D2-MSNs in stress-related outcomes.

## Material and methods

### Animals

All animal experimental protocols followed in this study were approved by the Institutional Animal Care and Use Committee of the University of South Carolina and the University of Maryland, Baltimore. Both female and male mice were used in the study, and all mice were maintained in a facility with a 12:12 reverse light-dark cycle (lights off at 7:00; lights on at 19:00) and provided with ad libitum access to food and water. All the behavior experiments were performed during the active period of mice. Mice breeders were procured from Jackson Laboratory, and breeding colonies were maintained in the university animal facility. Tac1-IRES2-Cre mice (Jackson Laboratory strain 021877) were bred with tdTomato (Ai14, Jackson Laboratory strain 007914) reporter mice, generating Tac1-Cre/tdTomato reporter mouse line. To confirm the presence of Tac1-Cre and tdTomato allele, ear punch tissue samples were used for genotyping. Weaning was done at postnatal day 21, and same-sex housing groups were maintained. The colonies were housed in a climate-controlled facility.

### Pavlovian fear conditioning

Fear conditioning (FC) was conducted in sound-attenuating fear conditioning chambers (Med Associates, USA). Prior to each conditioning session, mice were habituated in a separate room for 30 min under the same lighting conditions used for that session (training or cue-recall). All the experimental mice were assigned to one of the four conditioning procedures: (a) Paired conditioning, Conditioned stimulus (CS+) paired with an unconditioned stimulus (US+) (CS+US+); (b) Tone only, CS+ with no unconditioned stimulus (US−) (CS+US−); (c) Shock only (US+), with no conditioned stimulus (CS−) (CS−US+); (d) Context only (CS−US−). Conditioning was conducted in context A for 1, 3, 5, or 7 days, during which mice were placed in a chamber with an exposed metal grid, white light, and a vanilla scent (10% vanilla extract in ethanol). Following a 2 min baseline, mice in the paired conditioning group (CS+US+) received a 10 sec white-noise cues (80 dB), which co-terminated with a 0.6 mA foot shock, separated by a 30 sec interval. Mice received two pairings each day. Control groups receive only the tone (CS+US−), shock (CS−US+), or context only (CS−US−). Post-conditioning, after 24 hrs, mice were placed in a new context (context B) to test cue recall, in which 5 white noise cues that did not produce extinction were provided at 30 sec intervals after a 2 min baseline. Context B contained a chamber with an opaque white plexiglass floor insert, a curved wall insert, near-infrared (NIR) light, and vinegar scent (1% acetic acid in water). Freezing behavior during conditioning and recall was analyzed and used as a metric for learned aversive behavior. Brains were collected immediately after the cue recall for whole-cell patch-clamp electrophysiology.

### Slice electrophysiology

Whole-cell electrophysiological recordings were performed on the cue recall day. Brains were extracted after brief 3% isoflurane anesthesia and cervical dislocation. 300 µm sections were cut on a vibratome (Leica VT1000S) in ice-cold N-Methyl D-Glucamine (NMDG) cutting artificial cerebrospinal fluid (ACSF) solution (in mM): 92 NMDG, 20 HEPES, 25 glucose, 30 sodium bicarbonate, 1.2 sodium phosphate, 2.5 potassium chloride, 5 sodium ascorbate, 3 sodium pyruvate, 2 thiourea, 10 magnesium, 0.5 calcium chloride, osmolarity 305–310 mOsm, pH 7.34. Immediately after slicing, slices were placed in warm (32 ºC) NMDG solution for 2–5 min, then transferred to HEPES holding solution at room temperature for 1 h, which was the same as NMDG solution except NMDG was replaced with 92 mM sodium chloride and contained 1 mM magnesium chloride and 2 mM calcium chloride. Recording was performed at 33ºC, and slices were washed at a rate of 2 mL/min with ACSF (in mM): 125 sodium chloride, 26 sodium bicarbonate, and 11 glucose, 2.5 potassium chloride, 1.25 sodium phosphate, 2.4 calcium chloride, 1 magnesium chloride, osmolarity 305– 310 mOsm, pH 7.34. To block inhibitory currents, synaptic recordings were performed by holding cells at −70 mV in the presence of picrotoxin (100 μM) added to the ACSF. Cesium methanesulfate with neurobiotin (0.1%) was used as the internal solution (in mM): 117 cesium methanesulfonate, 20 HEPES, 2.8 sodium chloride, 4 magnesium ATP, 0.3 sodium GTP, 0.4 EGTA, pH 7.30, osmolarity 280–287 mOsm.

Whole-cell recordings were obtained in the NAc core using borosilicate glass pipettes (1.5-3.0 MΩ). D2-MSNs were distinguished from D1-MSNs by the absence of Tdtomato fluorescence using an Olympus BX51WI microscope and an LED light source (Olympus). After giga-seal formation and breaking in the cell, cells were allowed to stabilize for 5 min before data acquisition. Signals were amplified using a Multiclamp 700B amplifier and digitized (Digidata 1550B, Molecular Devices) at a 10 kHz sampling rate and filtered at 1 kHz for spontaneous excitatory postsynaptic currents (sEPSCs) and 2-4 kHz for electrically evoked currents. Synaptic currents were evoked using a concentric bipolar stimulating electrode. AMPAR/NMDAR ratios were acquired by taking the peak of the inward current at −70 mV divided by the +40 mV outward current 50 msec after the start of the AMPAR response. Paired pulse ratios (PPRs) were acquired at a 50 msec interval and were calculated as response 2/response 1. All evoked currents were the average of at least 7 sweeps. Cells with <20MΩ series resistance were included in the analysis and cells showing 20% or greater change in series resistance were excluded from analysis. Following recording, slices were collected in a 4% paraformaldehyde solution in 1X PBS for 24hrs before being placed in 1X PBS solution in 0.01% sodium azide for processing.

### Neuronal morphology

Neurons filled with neurobiotin were processed for neuronal morphology assessment. PFA-fixed slices in 0.01% sodium azide were washed in 1X PBS three times for 30 min each. Sections were then incubated in a blocking solution of 1X PBS with 0.5% Triton X-100 (20%), 10% Normal Donkey Serum (NDS) for 5 hrs at room temperature. Slices were then incubated overnight at room temperature in 1:1000 Steptavadin-488 (Jackson Laboratory, cat#016-540-084) in a modified blocking solution with 0.05% Triton X-100 (20%) and 10% NDS in 1XP BS. To remove unbound Streptavidin-488, slices were washed 3 times in 1X PBS for 30 minutes each. Slices were placed in 1X PBS, carefully mounted, dried, and cover-slipped using DAPI mounting medium. All images were acquired using a Leica TCS SP8 multiphoton confocal microscopy system (Leica Microsystems, Germany) and LASX software (Leica Microsystems, Germany). The images were captured at 40x-63x magnification using a GFP filter to visualize the streptavidin-488 tagged neurobiotin-filled neuron. Secondary dendrites were identified and imaged using Leica’s z-stack function to maintain a 3-D information of spines (see **Supplemental Figure 3**).

Spines were quantified using ImageJ. Secondary dendrites 10-30 µm in length and perpendicular to the beam path were chosen for quantification. Spines were quantified in each plane of the z-stack without using a z-stack projection. Spine types (Mushroom, Stubby, and Thin) were empirically determined using previous and current hand-scored datasets, which quantified head size and neck length and identified spine types by eye. Spine types were assigned using the following parameters: spine head area (µm^2^), neck length (µm)): Mushroom (≥0.2, >0.2); Stubby (>0.2, <0.2); Thin (<0.2, >0.2).

### Statistical analysis

Detailed statistics are provided in **Supplementary Table 1**. The reported data were analyzed, and statistics were calculated using GraphPad Prism software, and significance was considered if *P*<0.05. For all multi-group data, two-way ANOVAs were used followed by post-hoc tests using Tukey’s multiple comparison test. Student’s *t*-test was used only for the L-733,060 and saline injection groups in the 7-day CS+US+ condition. All data are represented as the mean ± SEM.

## Results

### Repeated fear conditioning increases cue-dependent learning

Our previous work and others have indicated that NAc D2-MSN activation facilitates social avoidance learning, but not depression-related outcomes and D2-MSN activity is driven by increased spine density ^8,26^. To determine if repeated conditioning produces the same D2-MSN effects as repeated social stress, Tac1-Cre/tdTomato mice were subjected to a Pavlovian fear conditioning protocol and underwent 1, 3, 5, and 7 days of conditioning (**Fig. 1a**). Following 1-day conditioning, freezing was tested in a novel context to determine the extent to which mice associated the cue with the aversive foot shock. As we have shown previously ^10^, two pairings of conditioned stimulus (CS) paired with an unconditioned stimulus (US) foot shock (CS+US+), led to significant freezing, compared to unpaired groups (CS−US−, CS−US+, CS+US−) (**Fig. 1b**), confirming cue-dependent learning. In addition, mice showed an enhancement in freezing to the cue as well as the conditioning context. After 7 days of repeated conditioning, freezing during recall was higher in the CS+US+ group than other groups (**Fig. 1c**). Baseline freezing was similar across all groups, with moderate freezing observed in the US+ condition for both CS− and CS+ groups, consistent with stress sensitization or context generalization ^27,28^. As expected, repeated conditioning days augmented freezing to both the context and the tone (**Fig. 1d**).

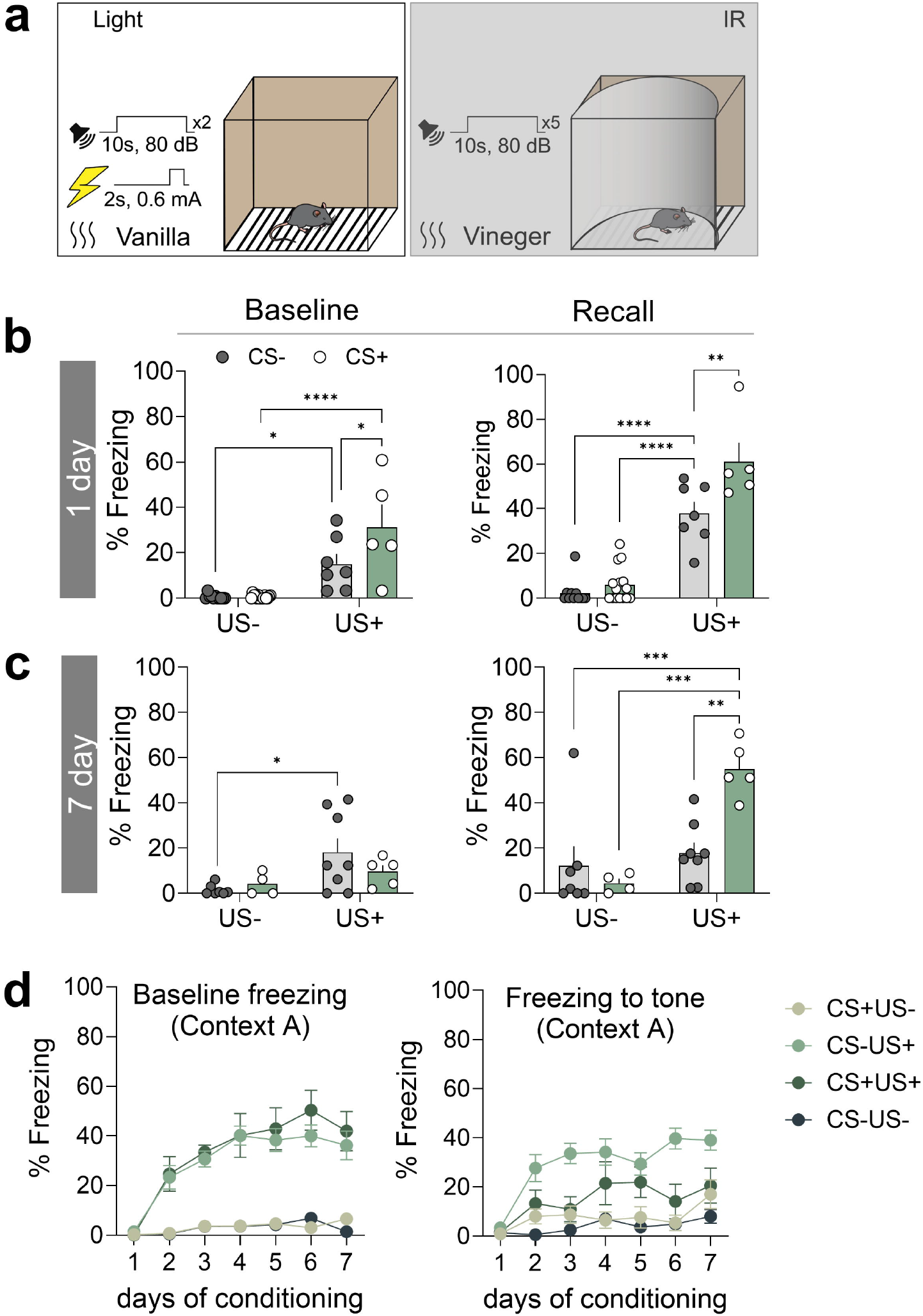

### Repeated fear conditioning alters synaptic inputs

To understand the underlying synaptic changes in D2-MSNs involved in the cue-dependent learning for aversive cues, we performed whole-cell patch clamp recordings from D2-MSNs (**Fig. 2**). Following a single day of conditioning, spontaneous excitatory postsynaptic current (sEPSC) amplitude was increased in D2-MSNs in the CS+/US+ group (**Fig. 2a**), as shown previously ^10^. sEPSC frequency, AMPAR/NMDAR ratios, and PPR showed no change across all other conditions (**Fig. 2a-b**). In the 3- and 5-day conditioning groups, no differences were observed (**Supplementary Fig. 1**). In contrast, 7-day conditioning animals revealed a selective increase in frequency in the CS+US+ group compared to the CS−US−group, with no change in CS conditions (**Fig. 2c**). These higher rates of spontaneous excitatory events, accompanied by no change in amplitude, AMPAR/NMDAR ratios, or PPR (**Fig. 2c-d**), and in the rectification **(Supplementary Fig. 2)**, implicating a shift in plasticity that may underlie long-term learning.

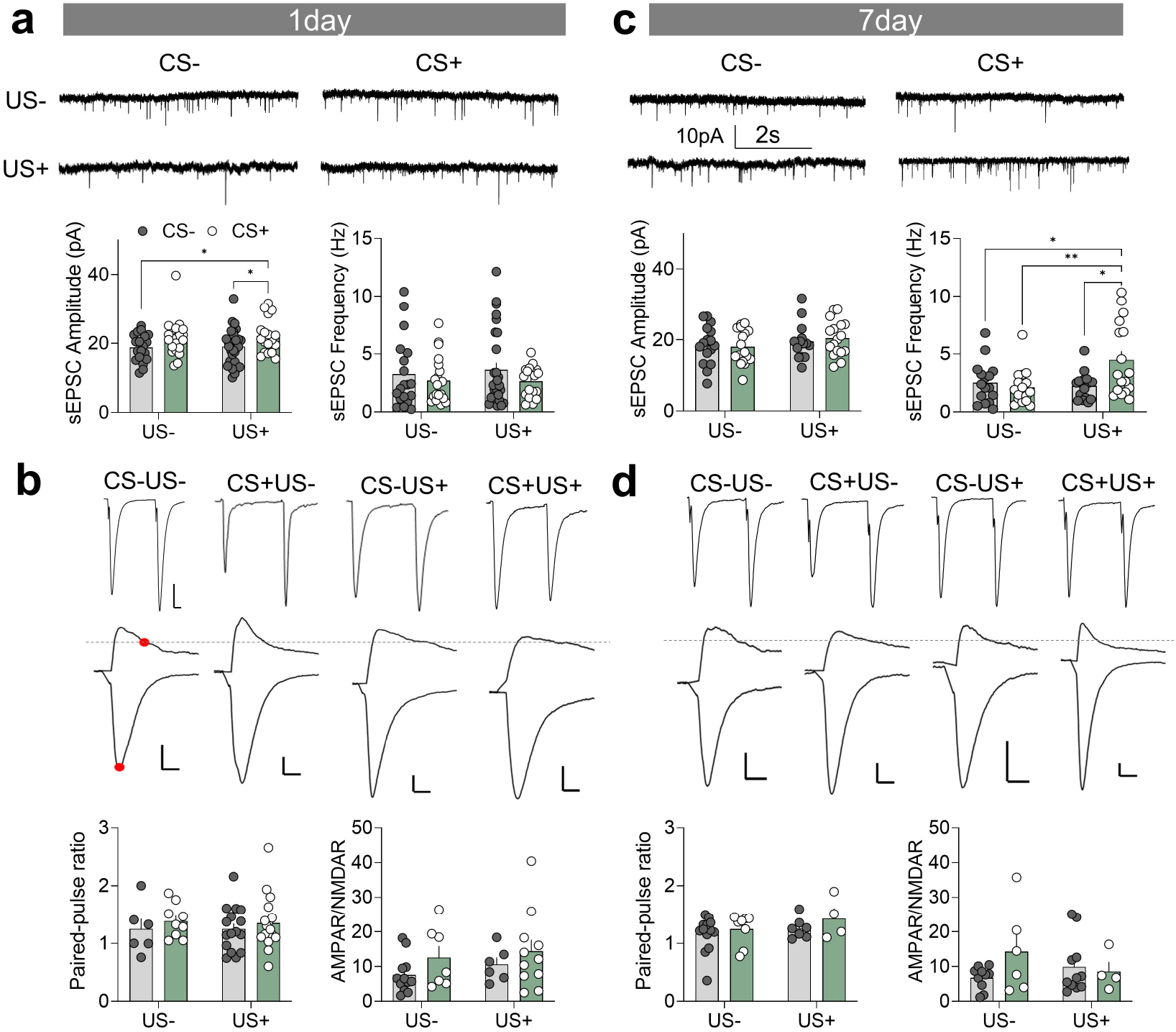

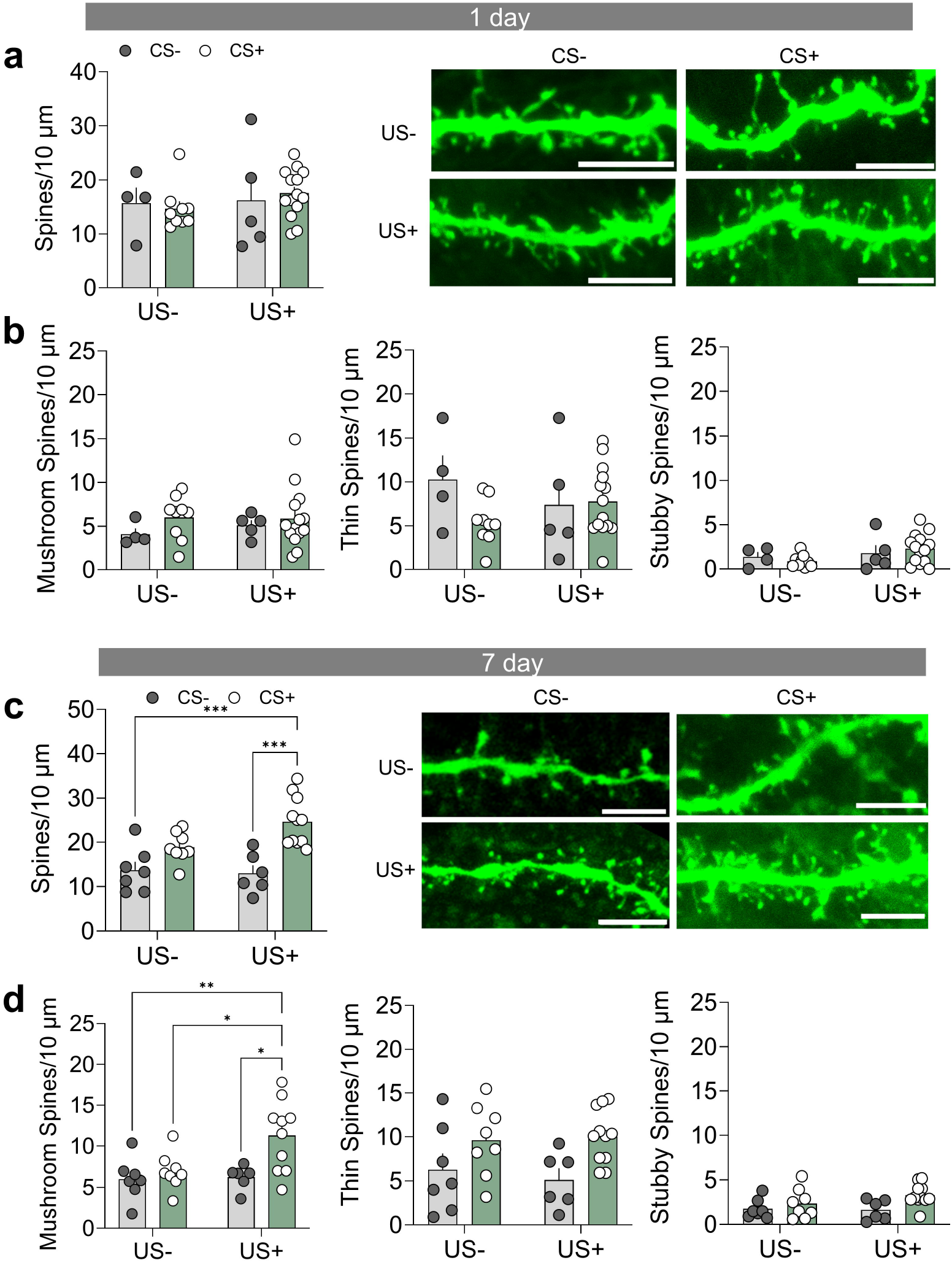

### Repeated conditioning shifts plasticity by increasing the number of spines

Increased sEPSC frequency, without a change in PPR, indicates increased excitatory connectivity onto D2-MSNs which could be mediated by an increase in spine density. To directly test whether this is due to structural synaptic remodeling and link it to freezing behavior, dendritic spine density and morphology were quantified in the same recorded neurons (**Fig. 3; Supplementary Fig. 3**). After one day of conditioning, no differences were observed in spine density (**Fig 3a-b**). However, after 7 days of repeated conditioning, the total spine density was significantly increased in CS+US+ group compared to all other groups (**Fig. 3c-d**). Furthermore, classification of spine subtypes shows an increase in thin and mushroom spines in the CS+US+ group ^29^, and no change in stubby spines (**Fig. 3d**). These results suggest spine formation supports the long-term cue-dependent learning observed in 7-day freezing and may underlie the increase in frequency of excitatory transmission.

### Blocking substance P signaling reduces release probability

The striatum has dense expression of the primary receptor for substance P, the neurokinin 1 receptor (NK1R) ^30^, and activation of the NK1R promotes selective excitatory plasticity on NAc D2-MSNs ^31^. Our previous work has shown that NAc substance P signaling is required for cue-dependent aversive learning from one day of fear conditioning ^10^. In addition, systemic and local antagonism or NAc knockout of the NK1R prior to one day conditioning prevents enhanced synaptic strength on D2-MSNs ^10,30^. Therefore, we tested whether repeated antagonism of the NK1R could block physiological changes caused by repeated conditioning. L-733,060 selectively increased PPR without affecting sEPSC amplitude or frequency (**Fig. 4a-b**) or AMPAR/NMDAR ratios (**Fig. 4c**), indicating reduced presynaptic release probability. This suggests an unknown and putatively active mechanism to blunt fear memories, caused by antagonism of the NK1R.

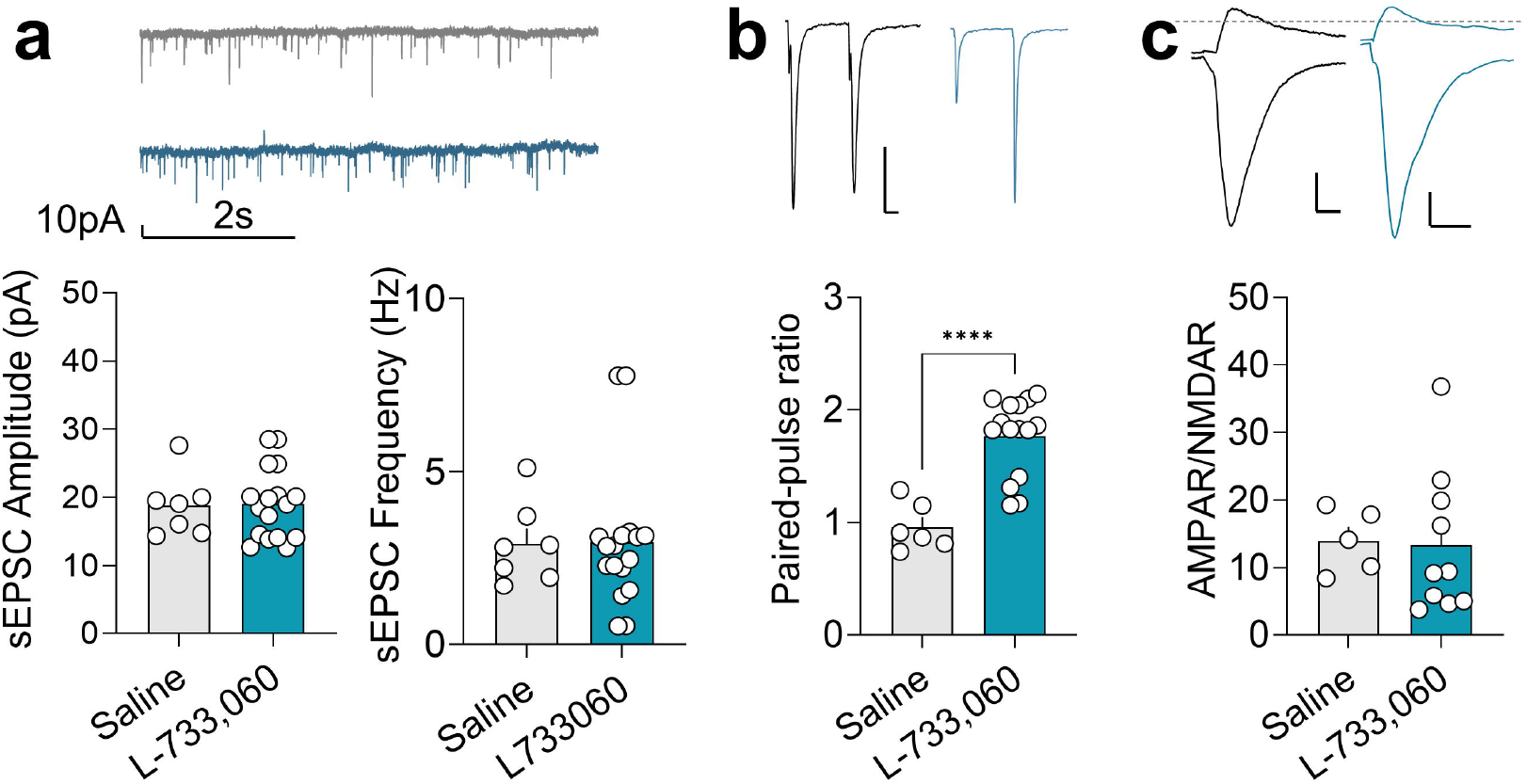

## Discussion

In this study, our data verify our previous results that NAc D2-MSN plasticity is caused by cue-dependent conditioning ^10^ and extends this dataset to show that repeated conditioning leads to significant and, sustained changes in plasticity through an increase in spine density. Furthermore, our data indicates that conditioned cues are required for this plasticity, not just foot shock alone, and this plasticity can be blunted by additional mechanisms to reduce excitatory transmission on NAc D2-MSNs. These results provide evidence that learned aversive stimuli, not just the aversive, stressful stimuli alone, promote D2-MSN plasticity. Previous studies have shown that repeated stress increases the density of excitatory spines on D2-MSNs ^21^. Electrophysiological measures recorded from 7 days CS+US+ conditioning mice showed that the postsynaptic excitatory input on NAc D2-MSNs is increased via an increase in excitatory spine density sEPSCs. This outcome is similar, if not equivalent, with earlier studies where mice received chronic stress ^8,32^. Furthermore, activation of NAc D2-MSNs specifically during an aversive social cue produces enhanced social avoidance without explicitly causing anhedonia ^8^. Concurrently, spine alterations, which are an important hallmark of dynamic neuronal plasticity and are important in learning ^33^, were not observed in cue-independent contextual learning following foot shock only. This supports our idea that structural remodeling at these synapses is associated with the formation of a stabilized fear memory that is cue-selective and not necessarily a generalized response to aversive cues. Together, these data suggest D2-MSN synapses are engaged in cue-dependent fear learning through coordinated structural and functional mechanisms.

Substance P is highly expressed in NAc and acts via NK1R ^10,30,31,34^, and its signaling is known to be important for learning about aversive cues ^12^. Interestingly, mice injected with NK1R antagonist allowed us to delineate learning-related synaptic plasticity from presynaptic modulation in NAc D2-MSNs. We found PPR was selectively increased at D2-MSNs synapses, indicating reduced presynaptic release probability that may compensate for increased frequency of excitatory input on D2-MSNs. We also observed no change in sEPSC amplitude and frequency, further revealing that this manipulation does not affect the exact mechanisms causing an increase in excitation that support learned behavior. These observations show that D2-MSNs fine-tune the circuit dynamics, consistent with a gain-setting modulatory role for substance P ^35^.

Taken together, our findings support a model **(Fig. 5**) in which fear learning is encoded through specific synaptic modifications on D2-MSNs that dissociate learned threat from generalized stress responses important for cue-dependent aversive response. We propose that the learning component of aversive conditioning is encoded by D2-MSNs and may contribute to how stress responses are perceived and processed.

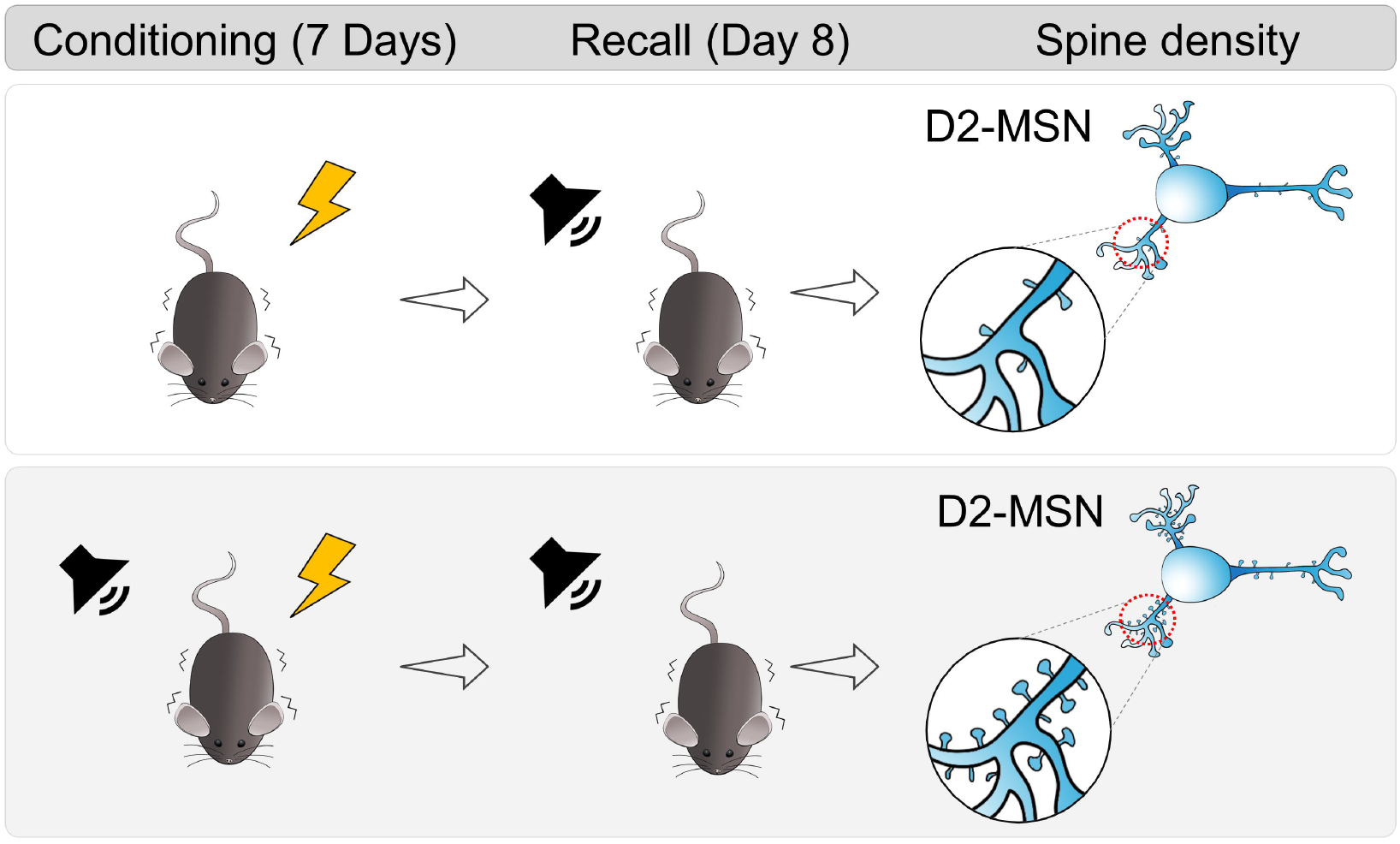

## Acknowledgements

This work was supported by NIH/NIMH R00MH123673, the Department of Drug Discovery and Biomedical Sciences in the College of Pharmacy at the University of South Carolina, and the Department of Psychiatry at the University of Maryland School of Medicine.

## Author Contributions

Conceptualization, D.D. and T.C.F.; methodology and investigation, D.D., C.G., E.L., H.K., M.H., and T.C.F.; writing, D.D., C.G., and T.C.F.; review and editing, D.D., E.L., and T.C.F.; supervision, T.C.F.

## Data Availability

Data is available upon request from the corresponding author.

## Additional Information

The authors declare no competing interests.

## References

1 James, P. H. et al. Regulation of the hypothalamic-pituitary-adrenocortical stress response. Compr Physiol 6, 603–621, (2016). 10.1002/cphy.c150015

2 Kaileigh, A. B. et al. Acute stress enhances tolerance of uncertainty during decision-making. Cognition 205 (2020). 10.1016/j.cognition.2020.104448

3 Gjerstad, J. K., Lightman, S. L. & Spiga, F. Role of glucocorticoid negative feedback in the regulation of HPA axis pulsatility. Stress 21, 403–416 (2018). 10.1080/10253890.2018.1470238

4 Ronald, S. D., Gerard, S. & John, H. K. Altered connectivity in depression: GABA and glutamate neurotransmitter deficits and reversal by novel treatments. Neuron 102, 75–90 (2019). 10.1016/j.neuron.2019.03.013

5 Francis, T. C. et al. Molecular basis of dendritic atrophy and activity in stress susceptibility. Mol Psychiatry 22, 1512–1519 (2017). 10.1038/mp.2017.178

6 Carlo, F. et al. Childhood stressful events, HPA axis and anxiety disorders. World J Psychiatry 2, 13–25, (2012). 10.5498/wjp.v2.i1.13

7 Sameer, H. Chronic stress, neuroinflammation, and depression: an overview of pathophysiological mechanisms and emerging anti-inflammatories. Front Psychiatry 14 (2023). 10.3389/fpsyt.2023.1130989

8 Francis, T. C. et al. Nucleus accumbens medium spiny neuron subtypes mediate depression-related outcomes to social defeat stress. Biol Psychiatry 77, 212–222 (2015). 10.1016/j.biopsych.2014.07.021

9 Francis, T. C. & Lobo, M. K. Emerging role for nucleus accumbens medium spiny neuron subtypes in depression. Biol Psychiatry 81, 645–653 (2017). 10.1016/j.biopsych.2016.09.007

10 Belilos, A. et al. Nucleus accumbens local circuit for cue-dependent aversive learning. Cell Rep 42, 113488 (2023). 10.1016/j.celrep.2023.113488

11 Carina, S.-C., Barbara, C., Nuno, S. & Ana, J. R. Reappraising striatal D1- and D2-neurons in reward and aversion. Neurosci Biobehav Rev 68, 370–386, Elsevier (2016). 10.1016/j.neubiorev.2016.05.021

12 Jennifer, E. Z. et al. D1 and D2 medium spiny neurons in the nucleus accumbens core have distinct and valence-independent roles in learning. Neuron 112, 835–849.e837, (2024). 10.1016/j.neuron.2023.11.023

13 You Hsin, L. et al. Accumbal D2R-medium spiny neurons regulate aversive behaviors through PKA-Rap1 pathway. Neurochem Int 143 (2021). 10.1016/j.neuint.2020.104935

14 Carina, S.-C. et al. Nucleus accumbens medium spiny neurons subtypes signal both reward and aversion. Mol Psychiatry 25, 3241–3255, Mol Psychiatry (2020). 10.1038/s41380-019-0484-3

15 Elena Chaves, R., Jérémie, N., Daniel, R. & Alban de, K. Direct and indirect striatal projecting neurons exert strategy-dependent effects on decision-making. (2025).

16 Jérôme, L., Alexander, S. J., Colin, M. & Jentsch, J. D. Dopamine D2 receptors in dopaminergic neurons modulate performance in a reversal learning task in mice. eNeuro 5 (2018). 10.1523/ENEURO.0229-17.2018

17 Tom, M., Ji Yoon, K. & Takatoshi, H. Nucleus accumbens core dopamine D2 receptor-expressing neurons control reversal learning but not set-shifting in behavioral flexibility in male mice. Front Neurosci 16 (2022). 10.3389/fnins.2022.885380

18 Yudan, D. et al. Reduced nucleus accumbens functional connectivity in reward network and default mode network in patients with recurrent major depressive disorder. Transl Psychiatry 12 (2022). 10.1038/s41398-022-01995-x

19 Bingqian, Z. et al. Alterations of static and dynamic functional connectivity of the nucleus accumbens in patients with major depressive disorder. Frontiers in Psychiatry 13 (2022). 10.3389/fpsyt.2022.877417

20 Daniel, J. C. et al. IκB kinase regulates social defeat stress-induced synaptic and behavioral plasticity. J Neurosci 31, 314–321 (2011). 10.1523/JNEUROSCI.4763-10.2011

21 Megan, E. F., Antonio, F., Miriam, S. M. & Mary Kay, L. Dendritic spine density is increased on nucleus accumbens D2 neurons after chronic social defeat. Sci. Rep. 10 (2020). 10.1038/s41598-020-69339-7

22 Francis, T. C., Gaynor, A., Chandra, R., Fox, M. E. & Lobo, M. K. The selective RhoA inhibitor rhosin promotes stress resiliency through enhancing D1-medium spiny neuron plasticity and reducing hyperexcitability. Biol Psychiatry 85, 1001–1010 (2019). 10.1016/j.biopsych.2019.02.007

23 Lucantonio, F. et al. Ketamine rescues anhedonia by cell-type- and input-specific adaptations in the nucleus accumbens. Neuron 113, 1398–1412 e1394 (2025). 10.1016/j.neuron.2025.02.021

24 Pignatelli, M. et al. Cooperative synaptic and intrinsic plasticity in a disynaptic limbic circuit drive stress-induced anhedonia and passive coping in mice. Mol Psychiatry 26, 1860–1879 (2021). 10.1038/s41380-020-0686-8

25 Lim, B. K., Huang, K. W., Grueter, B. A., Rothwell, P. E. & Malenka, R. C. Anhedonia requires MC4R-mediated synaptic adaptations in nucleus accumbens. Nature 487, 183–189 (2012). 10.1038/nature11160

26 Fox, M. & Lobo, M. The molecular and cellular mechanisms of depression: a focus on reward circuitry. Mol Psychiatr, 1–18 (2019). 10.1038/s41380-019-0415-3

27 Arp, J. M. et al. Blocking glucocorticoid receptors at adolescent age prevents enhanced freezing between repeated cue-exposures after conditioned fear in adult mice raised under chronic early life stress. Neurobiol Learn Mem 133, 30–38, (2016). 10.1016/j.nlm.2016.05.009

28 Jiahui, Y., Toshie, N. & Masanori, S. Fear generalization immediately after contextual fear memory consolidation in mice. Biochem Biophys Res Commun 558, 102–106, (2021). 10.1016/j.bbrc.2021.04.072

29 Kalen, P. B. & Elly, N. Spine dynamics: are they all the same? Neuron 96, 43–55 (2017). 10.1016/j.neuron.2017.08.008

30 Virginia, M. P., Jennifer, D., June, C., Patrick, D. G. & Nigel, W. B. Neurokinin 1 receptor distribution in cholinergic neurons and targets of substance P terminals in the rat nucleus accumbens. J Comp Neurol 423, 500–511 (2000). 10.1002/1096-9861(20000731)423:33.0.CO;2-9

31 Francis, T. C., Yano, H., Demarest, T. G., Shen, H. & Bonci, A. High-frequency activation of nucleus accumbens D1-msns drives excitatory potentiation on D2-msns. Neuron 103, 432–444 e433 (2019). 10.1016/j.neuron.2019.05.031

32 Lena, A. K. et al. Stress and Cocaine Trigger Divergent and Cell Type. 2013;Specific Regulation of Synaptic Transmission at Single Spines in Nucleus Accumbens. Biol Psychiatry 79, 898–905, Elsevier (2016). 10.1016/j.biopsych.2015.05.022

33 Gipson, C. D. & Olive, M. F. Structural and functional plasticity of dendritic spines - root or result of behavior? Genes Brain Behav 16, 101–117 (2017). 10.1111/gbb.12324

34 Toshihiko, A. & Yasuo, K. Actions of substance P on rat neostriatal neurons in vitro. The Journal of Neuroscience 16, 5141–5153 (1996). 10.1523/JNEUROSCI.16-16-05141.1996,

35 Zi Xuan, H., Ting Yu, L., Yue Yue, Y., Hui Fang, S. & Xiao Juan, Z. Substance P plays a critical role in synaptic transmission in striatal neurons. Biochem Biophys Res Commun 511, 369–373, Elsevier B.V. (2019). 10.1016/j.bbrc.2019.02.055

