## Supplementary Information for "Cue-Dependent Fear Learning Drives Nucleus Accumbens Spine Plasticity"

### Supplementary Figures and Tables

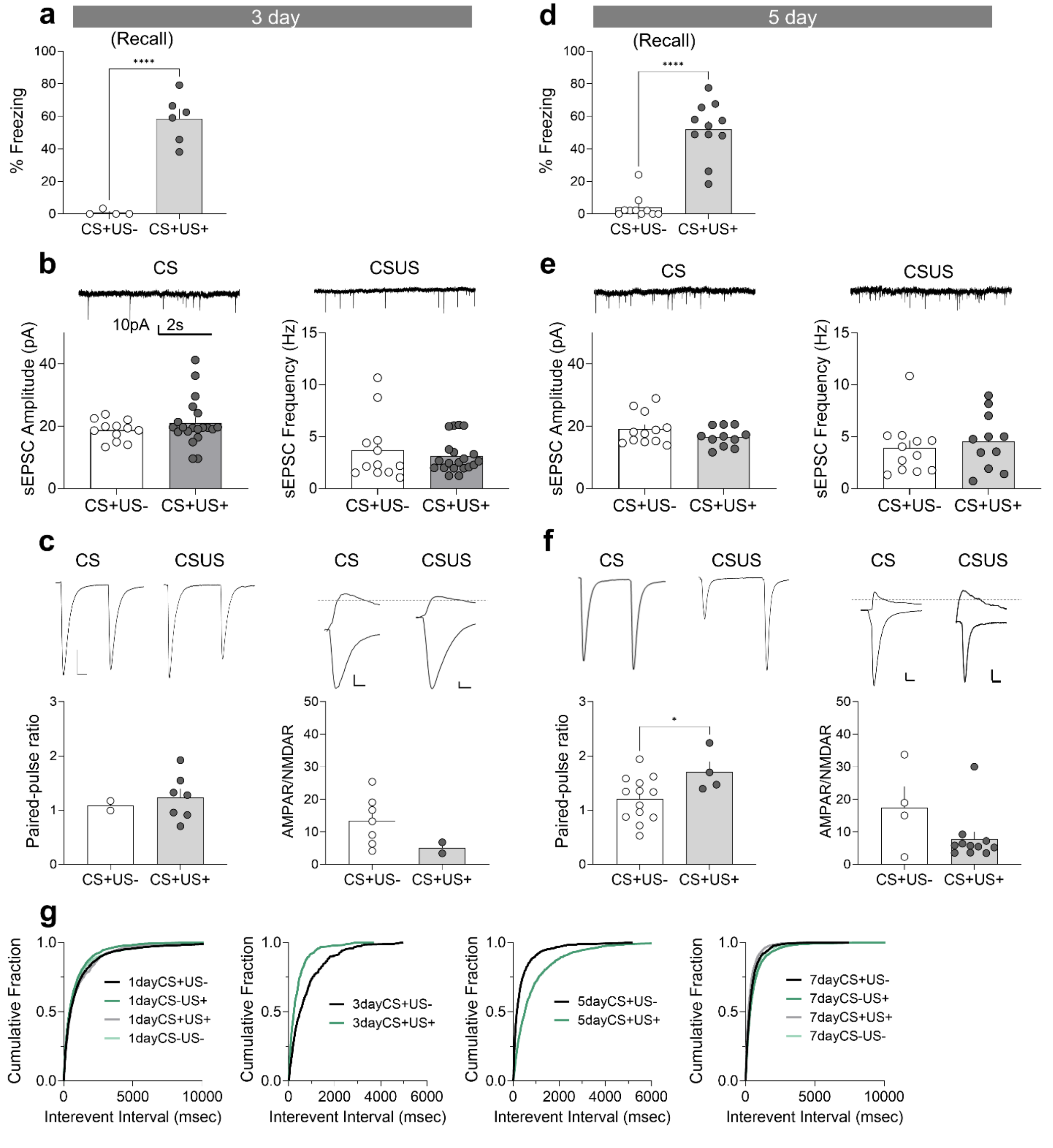


**Supplementary Figure 1. Electrophysiological measures are unchanged following 3 or 5 days of conditioning. (a)** 3-days of conditioning enhanced freezing to the cue ($t(8)=7.634$, $P<0.0001$). **(b)** sEPSC amplitude and frequency remained unchanged after 3-days of conditioning. Mice/cells (N/n): CS+US-=4/12, CS+US+=6/20. **(c)** PPR, and AMPAR/NMDAR ratios remained unchanged after 3-days of conditioning.

**(d)** 5-days of conditioning enhanced freezing to the cue ($t(19) = 8.087$ $P<0.0001$). N/n: CS+US-=4/2, CS+US+=6/7. **(e)** sEPSCs amplitude and frequency and AMPAR/NMDAR ratios remained unchanged after 5-days of conditioning. N/n: CS+US-=10/12, CS+US+=11/11. **(f)** PPR was changed after 5-days of conditioning ($P<0.0496$, $t(15) = 2.136$) while AMPAR/NMDAR ratios remained unchanged. N/n: CS+US-=10/13, CS+US+=11/4. **(g)** Cumulative distribution of sEPSC interevent intervals across experimental days and with different conditions. Scale (PPR: 100pA/10msec, ampar/nmdar: 50pA/50msec). Data are represented as mean± SEM, *$P<0.05$, ****$P<0.0001$.

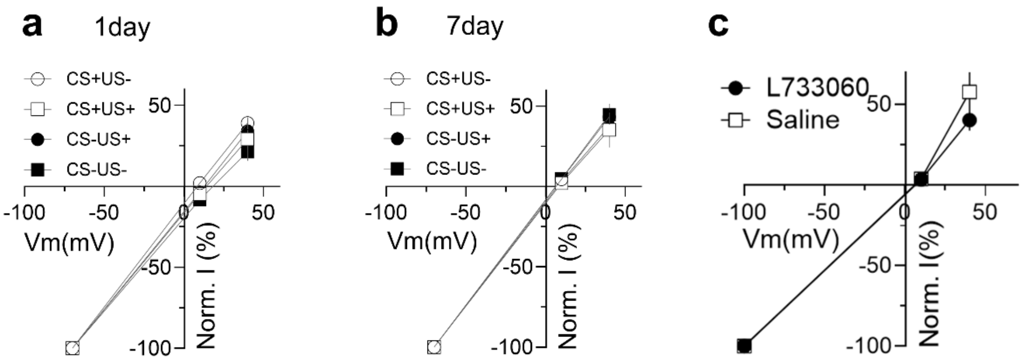


**Supplementary Figure 2. AMPAR IVs remain unchanged after fear conditioning.** AMPAR-evoked current-voltage relationship remained unchanged after **(a)** 1-day (Two-way RM ANOVA, no interaction, *F*(6,82) =0.4493, *P*=0.8435*,* N/n: CS+US-=5/8, CS+US+= 7/13, CS-US+= 7/16, CS-US-= 3/8) or **(b)** 7-days (Two-way RM ANOVA, no interaction, *F*(6,38) =0.3812, *P*=0.8864*,* N/n: CS+US-= 3/5, CS+US+= 1/2, CS-US+= 5/8, CS-US-= 4/8) of conditioning. **(c)** No change after 7-days of conditioning was observed in AMPAR IV curves in Sal or L-733,060 injected mice (Two-way RM ANOVA, no interaction, *F*(2, 18) =0.9482, *P=0.4059*). N/n: Sal= 5/7, L-733060= 3/4. Data is represented as mean± SEM.

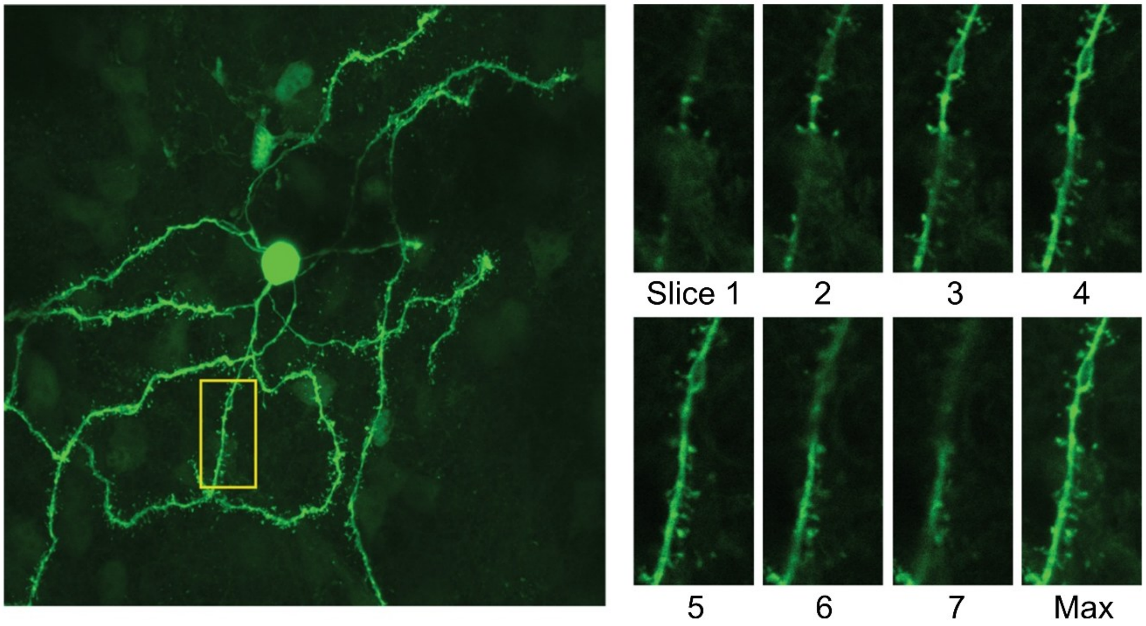


**Supplementary Figure 3. Three-dimensional spine analysis.** (*left*) Images of dendritic arbors were acquired and secondary dendrites within the plane of the focus were isolated and analyzed. (*right*) An individual secondary spine from the yellow insert. Dendritic spines were quantified using individual z-slices. Three-dimensional images were used to decrease loss of information by a stack maximum (Max).

### Supplementary Table 1: Exact Statistics

| Fig | Panel | Test | Test Statistic | | *P*-value | N |
|---|---|---|---|---|---|---|
| 1 | (b) Baseline freezing | Two-way ANOVA | CS, US | Interaction $F(1, 34)$ =6.281 | 0.0172 | 11,15,7,5 |
| | | | | Main effect of US, $F(1,34)$ =49.01 | 0.0001 | |
| | | | | Main effect of CS, $F(1, 34)$ =6.408 | 0.0162 | |
| | | | CS-US- vs CS+US- | Tukey's mc $q(34)$ =0.0317 | 0.9999 | |
| | | | CS-US- vs CS-US+ | Tukey's mc $q(34)$ =4.650 | 0.0120 | |
| | | | CS-US- vs CS+US+ | Tukey's mc $q(34)$ =8.840 | 0.0001 | |
| | | | CS+US- vs CS-US+ | Tukey's mc q(34) =4.884 | 0.0078 | |
| | | | CS+US- vs CS+US+ | Tukey's mc $q(34)$ =9.209 | 0.0001 | |
| | | | CS-US+ vs CS+US+ | Tukey's mc $q(34)$ =4.303 | 0.0222 | |
| | Recall freezing | Two-way ANOVA | CS, US | Interaction $F(1,34)$ =6.971 | 0.0124 | |
| | | | | Main effect of CS, $F(1,34)$ =13.03 | 0.0010 | |
| | | | | Main effect of US, $F(1,34)$ =148.5 | 0.0001 | |
| | | | CS-US- vs CS-US- | Tukey's mc $q(34)$ =1.222 | 0.8233 | |
| | | | CS-US- vs CS-US+ | Tukey's mc $q(34)$ =9.875 | 0.0001 | |
| | | | CS-US- vs CS+US+ | Tukey's mc $q(34)$ =14.65 | 0.0001 | |
| | | | CS+US- vs CS-US+ | Tukey's mc $q(34)$ =9.371 | 0.0001 | |
| | | | CS+US- vs CS+US+ | Tukey's mc $q(34)$ =14.36 | 0.0001 | |
| | | | CS-US+ vs CS+US+ | Tukey's mc $q(34)$ = 5.339 | 0.0033 | |
| | (c) Baseline freezing | Two-way ANOVA | CS, US | No interaction $F(1,20)$ =1.469 | 0.2396 | 7,4,8,5 |
| | | | | Main effect of CS, $F(1,20)$ =0.416 | 0.5263 | |
| | | | | Main effect of US, $F(1,20)$ =5.805 | 0.0257 | |
| | | Two-way ANOVA | CS, US | Interaction $F(1,20)$ =11.45 | 0.0029 | |
| | | | | Main effect of CS, $F(1,20)$ =4.870 | 0.0392 | |
| | | | | Main effect of US, $F(1,20)$ =17.74 | 0.0004 | |
| | | | CS-US- vs CS+US- | Tukey's mc $q(20)$ =1.125 | 0.8556 | |
| | | | CS-US- vs CS-US+ | Tukey's mc $q(20)$ =0.98581 | 0.9044 | |
| | | | CS-US- vs CS+US+ | Tukey's mc $q(20)$ =6.567 | 0.0008 | |
| | | | CS+US- vs CS-US+ | Tukey's mc $q(20)$ =1.961 | 0.5215 | |
| | | | CS+US- vs CS+US+ | Tukey's mc $q(20)$ =6.783 | 0.0006 | |
| | | | CS-US+ vs CS+US+ | Tukey's mc $q(20)$ =5.875 | 0.0025 | |
| 2 | (a) sEPSC amplitude | Two-way ANOVA | CS, US | No interaction $F(1, 82)$ =0.014 | 0.9055 | 19,21,28,18 |
| | | | | Main effect of CS, $F(1,82)$ =7.304 | 0.0084 | |

| | | | | | | |
|---|---|---|---|---|---|---|
| | | | | Main effect of US, $F(1,82)$ =0.1313 | 0.7181 | |
| | sEPSC frequency | Two-way ANOVA | CS, US | No interaction $F(1, 82)$ =0.1353<br>Main effect of CS, $F(1,82)$ =1.830<br>Main effect of US, $F(1,82)$ =0.07 | 0.7139<br>0.1798<br>0.792 | |
| | (b) sEPSC amplitude | Two-way ANOVA | CS, US | No interaction $F(1, 60)$ =0.3432<br>Main effect of CS, $F(1,60)$ =0.017<br>Main effect of US, $F(1,60)$ =1.722 | 0.5602<br>0.8942<br>0.1944 | 16,17,14,17 |
| | sEPSC frequency | Two-way ANOVA | CS, US | Interaction $F(1, 62)$ =6.799<br>Main effect of CS, $F(1,62)$ =3.284<br>Main effect of US, $F(1,62)$ =4.247 | 0.0114<br>0.0748<br>0.0435 | |
| | | | CS-US- vs CS+US- | Tukey's mc test, $q(62)$=0.7952 | 0.9428 | |
| | | | CS-US- vs CS+US- | Tukey's mc test, $q(62)$=0.5384 | 0.9810 | |
| | | | CS-US- vs CS+US- | Tukey's mc test, $q(62)$=3.873 | 0.0392 | |
| | | | CS-US- vs CS+US- | Tukey's mc test, $q(62)$=0.2487 | 0.9980 | |
| | | | CS-US- vs CS+US- | Tukey's mc test, $q(62)$=4.741 | 0.0073 | |
| | | | CS-US- vs CS+US- | Tukey's mc test, $q(62)$=4.420 | 0.0140 | |
| | (c)PPR | Two-way ANOVA | CS, US | Interaction $F(1, 44)$ =0.018<br>Main effect of CS, $F(1,44)$ =0.02<br>Main effect of US, $F(1,44)$ =0.8436 | 0.8926<br>0.3634<br>0.888 | 6,9,16,17 |
| | AMPAR/NMDAR | Two-way ANOVA | CS, US | Interaction $F(1, 32)$ =0.043<br>Main effect of CS, $F(1,32)$ =02.421<br>Main effect of US, $F(1,32)$ =0.7178 | 0.8365<br>0.4032<br>0.1296 | 12,7,6,11 |
| | (d) PPR | Two-way ANOVA | CS, US | Interaction $F(1, 28)$ =0.1503<br>Main effect of CS, $F(1,28)$ =1.350<br>Main effect of US, $F(1,28)$ =1.682 | 0.7011<br>0.2551<br>0.2052 | 13,8,7,4 |
| | AMPAR/NMDAR | Two-way ANOVA | CS, US | Interaction $F(1, 27)$ =0.2193<br>Main effect of CS, $F(1,27)$ =1.047<br>Main effect of US, $F(1,27)$ =0.1896 | 0.1502<br>0.3154<br>0.6667 | 10,6,11,4 |
| 3 | (a)total spines | Two-way ANOVA | CS, US | No interaction $F(1, 28)$ =0.2898<br>Main effect of CS, $F(1,28)$ =0.004<br>Main effect of US, $F(1,28)$ =0.5737 | 0.5946<br>0.9476<br>0.4551 | 4,9,5,14 |
| | Mushroom spines | Two-way ANOVA | CS, US | No interaction $F(1, 28)$ =0.2561<br>Main effect of CS, $F(1,28)$ =1.428<br>Main effect of US, $F(1,28)$ =0.1580 | 0.6168<br>0.2422<br>0.6940 | |
| | Thin spines | Two-way ANOVA | CS, US | No interaction $F(1, 28)$ =2.408<br>Main effect of CS, $F(1,28)$ =1.770<br>Main effect of US, $F(1,28)$ =0.023 | 0.1319<br>0.1941<br>0.8801 | |

| | | | | | | |
|---|---|---|---|---|---|---|
| | Stubby spines | Two-way ANOVA | CS, US | No interaction *F*(1, 28) =0.7226<br>Main effect of CS, *F*(1,28) =.0007<br>Main effect of US, *F*(1,28) =2.413 | 0.4025<br>0.9779<br>0.1316 | |
| | (c) total spines | Two-way ANOVA | CS, US | No interaction *F*(1, 28) =3.549<br>Main effect of CS, *F(1,28)* =24<br>Main effect of CS, *F(1,28)* =2.127 | 0.070<br>0.0001<br>0.1558 | 7,8,6,11 |
| | (d) mushroom spines | Two-way ANOVA | CS, US | No interaction *F*(1, 27) =3.442<br>Main effect of CS, *F*(1, 27) =7.261<br>Main effect of US, *F*(1, 27) = 4.238 | 0.0745<br>0.0120<br>0.0493 | |
| | | | CS-US- vs CS+US- | Tukey's mc q(28) =0.8384 | 0.9394 | |
| | | | CS-US- vs CS+US- | Tukey's mc q(28) =02.1889 | 0.9991 | |
| | | | CS-US- vs CS+US+ | Tukey's mc q(28) =4.986 | 0.0078 | |
| | | | CS-US- vs CS+US | Tukey's mc q(28) =0.6088 | 0.9727 | |
| | | | CS+US- vs CS+US+ | Tukey's mc q(28) = 4.266 | 0.0266 | |
| | | | CS-US+ vs CS+US+ | Tukey's mc q(28) =4.555 | 0.0165 | |
| | Thin spines | Two-way ANOVA | CS, US | No interaction *F*(1, 28) =0.3063<br>Main effect of CS, *F*(1,28) =9.010<br>Main effect of US, *F*(1,28) =0.6797 | 0.5844<br>0.0056<br>0.7962 | |
| | Stubby spines | Two-way ANOVA | CS, US | No interaction *F*(1, 28) =1.311<br>Main effect of CS, *F*(1,28) =5.624<br>Main effect of US, *F*(1,28) =0.7584 | 0.2618<br>0.0248<br>0.3912 | |
| 4 | (a) sEPSC amplitude | Two-tailed unpaired *t*-test | Saline vs L-733,060 | *t*(22)=0.1215 | 0.9044 | 7,17 |
| | sEPSC Frequency | Two-tailed unpaired *t*-test | Saline vs L-733,060 | *t*(22)=0.0562 | 0.9556 | |
| | (b) Paired-pulse ratio | Two-tailed unpaired *t*-test | Saline vs L-733,060 | *t*(19) =5.357 | 0.0001 | 6,15 |
| | (c)AMPAR/NMDAR | Two-tailed unpaired *t*-test | Saline vs L-733,060 | *t*(13) =0.1229 | 0.9041 | 5,10 |
